# Consistent measurement of LAG-3 expression across multiple staining platforms with the 17B4 antibody clone

**DOI:** 10.1101/2022.02.21.481075

**Authors:** John B. Wojcik, Keyur Desai, Konstantinos Avraam, Arno Vandebroek, Lloye M. Dillon, Giorgia Giacomazzi, Charlotte Rypens, Joseph L. Benci

**Author notes:** **Corresponding author:** Joseph L. Benci, Bristol Myers Squibb, Princeton, NJ 08540, USA.

## Abstract

**Context:** An immunohistochemistry (IHC) assay developed to detect lymphocyte-activation gene 3 (LAG-3), a novel immune checkpoint inhibitor target, has demonstrated high analytical precision and interlaboratory reproducibility using a Leica staining platform, but has not been investigated on other IHC staining platforms.

**Objective:** To evaluate the performance of LAG-3 IHC assays using the 17B4 antibody clone across widely used IHC staining platforms: Agilent/Dako Autostainer Link 48 (ASL-48) and VENTANA BenchMark ULTRA (VBU) compared with Leica BOND-RX (BOND-RX).

**Design:** Eighty formalin-fixed paraffin-embedded melanoma tissue blocks were cut into consecutive sections and evaluated using staining platform–specific IHC assays with the 17B4 antibody clone. Duplicate testing was performed on the BOND-RX platform to assess intraplatform agreement. LAG-3 expression using a numerical score was evaluated by a pathologist and with a digital scoring algorithm. LAG-3 positivity was determined from manual scores using a ≥ 1% cutoff.

**Results:** LAG-3 IHC staining patterns and intensities were visually similar across all 3 staining platforms. Pearson correlation was ≥ 0.88 for interplatform and BOND-RX intraplatform concordance when LAG-3 expression was evaluated with a numerical score determined by a pathologist. Correlation increased with a numerical score determined with a digital scoring algorithm (Pearson correlation ≥ 0.93 for all comparisons). Overall percentage agreement was ≥ 77.5% for interplatform and BOND-RX intraplatform comparisons when a ≥ 1% cutoff was used to determine LAG-3 positivity.

**Conclusions:** Data from this study demonstrate that LAG-3 expression can be robustly and reproducibly assessed across 3 major commercial IHC staining platforms using the 17B4 antibody clone.

## Introduction

Lymphocyte-activation gene 3 (LAG-3) is a cell-surface immune checkpoint molecule expressed on immune cells (ICs) initiating an inhibitory signal that can impair T-cell activity and attenuate proinflammatory cytokine responses.^1-3^ Preclinical studies have shown that dual blockade of LAG-3 and programmed death-1 (PD-1) has synergistic antitumor activity, indicating that LAG-3 is an ideal candidate for novel immune checkpoint inhibitor combinations.^4^ RELATIVITY-047 (NCT03470922), a phase 2/3, global, randomized, double-blind trial, evaluated combined LAG-3 and PD-1 inhibition with relatlimab, an anti–LAG-3 antibody, and nivolumab, an anti–PD-1 antibody, as a novel fixed-dose combination versus nivolumab in patients with previously untreated metastatic or unresectable melanoma.^5^ In this study, combined treatment with relatlimab and nivolumab demonstrated superior progression-free survival (PFS) compared with nivolumab monotherapy regardless of LAG-3 expression.^5^

LAG-3 expression may be reflective of tumor inflammation and therefore generally predictive of response to the immuno-oncology therapy class.^5-8^ Consequently, assessing LAG-3 expression by immunohistochemistry (IHC) remains of significant interest to the research community despite a lack of clinical utility in informing relatlimab treatment decisions in advanced melanoma.^5^ A LAG-3 IHC assay using the 17B4 LAG-3 antibody clone on the Leica BOND-III staining platform has been developed in collaboration between Bristol Myers Squibb and LabCorp.(Johnson et al., in preparation) This assay is currently being used in relatlimab clinical trials, including the recently completed RELATIVITY-047.^5^ Recent data demonstrated the reproducible intraobserver, interobserver, and interlaboratory performance of the LabCorp LAG-3 IHC assay using the 17B4 antibody clone on the Leica BOND-III platform but did not investigate its performance on other staining platforms.(Johnson et al., in preparation)

Although assessment of LAG-3 expression is not required in a diagnostic context, a crucial barrier to widespread implementation of an IHC assay is its ability to produce concordant results using different staining platforms.^5^ Commercial IHC assays that are developed for a specific staining platform can present a significant challenge to testing laboratories, which may not have compatible equipment. Laboratory developed tests enable the reliable use of an IHC assay on alternative staining platforms by adapting validated assays, but do not always produce concordant results. Stringent standardization is therefore recommended prior to routine clinical use.^9-11^ This highlights the importance of reliable cross-platform performance for IHC assays to enable broad implementation and maximum utility. The LAG-3 IHC assay described by Johnson et al. was developed using the 17B4 antibody clone on the Leica BOND-III platform.(Johnson et al, in preparation) However, other IHC staining platforms, such as the Agilent/Dako Autostainer Link 48 (ASL-48) and VENTANA BenchMark ULTRA (VBU) are also commonly used. In this study, we assessed the feasibility of developing staining procedures using the 17B4 antibody clone on other staining platforms. The aim of our study was to develop protocols using ASL-48 or VBU that produced results comparable to the Leica BOND-RX (BOND-RX) platform. Interplatform concordance was examined using both pathologist evaluation and an investigational digital pathology method.

## Methods

### Samples and staining procedures

Eighty formalin-fixed paraffin-embedded (FFPE) melanoma tissue blocks were obtained from commercial vendors (BioIVT, Detroit, MI, USA; Discovery Life Sciences, Huntsville, AL, USA). Each tissue block was cut at 4-μm thickness into 4 consecutive sections for LAG-3 IHC staining on each platform with additional sections cut for hematoxylin and eosin (H&E) staining and isotype controls. Consecutive sections were evaluated using BOND-RX (section 1), ASL-48 (section 2), VBU (section 3), and BOND-RX (section 4), with 2 sections assessed using BOND-RX to measure intraplatform agreement for evaluation (hereafter referred to as BOND-RX 1 [section 1] and BOND-RX 2 [section 4]) (**Figure 1**). Deparaffinization and melanin removal were performed as described by Johnson et al.(Johnson et al., in preparation) All experiments across all platforms were performed using a monoclonal LAG-3 antibody, clone 17B4 (Cat. # LS-C18692, LSBio, Seattle, WA). Staining procedures were initially developed on BOND-RX, after which protocols were iteratively optimized on ASL-48 and VBU to produce visually similar staining to BOND-RX, before performing comparisons. All staining was performed in a single laboratory to avoid interlaboratory bias. Staining procedures and reagents were specific to each platform and are summarized in **Table 1**. The study was performed in accordance with the Bristol Myers Squibb Bioethics policy (https://www.bms.com/about-us/responsibility/position-on-key-issues/bioethics-policy-statement.html) and adhered to the World Medical Association Declaration of Helsinki for Human Research.

**Figure 1.**
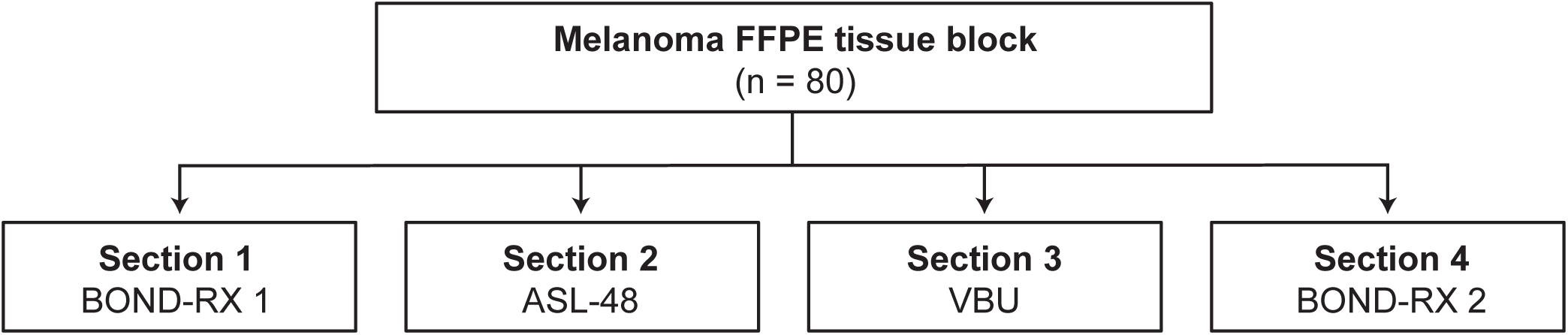
Diagram showing the consecutive sectioning design. Eighty commercially procured FFPE melanoma tissue blocks were cut into consecutive sections and stained using the LAG-3 17B4 antibody clone as shown. LAG-3 expression was analyzed with a 5-day washout period between sections corresponding to the same slide by a single pathologist. ASL-48, Agilent/Dako Autostainer Link 48; BOND-RX 1, Leica BOND-RX section 1; BOND-RX 2, Leica BOND-RX section 4; FFPE, formalin-fixed paraffin- embedded; LAG-3, lymphocyte-activation gene 3; VBU, VENTANA BenchMark ULTRA.

### Pathologist scoring

Slides were scored for LAG-3–positive IC content within the tumor region by a single expert pathologist trained on LAG-3 scoring methodology, as previously described.(Johnson et al, in preparation) Scoring was performed by a single pathologist to avoid interobserver bias. The tumor region included ≥ 100 tumor cells (TCs [confirmed using an H&E–stained slide]), intratumoral stroma, and peritumoral stroma (the band of stromal elements directly contiguous with the outer tumor margin) and did not include normal and/or adjacent uninvolved tissues. LAG-3 expression was recorded using a numerical score defined as the percentage of LAG-3–positive ICs that morphologically resembled lymphocytes relative to all nucleated cells (ICs [lymphocytes and macrophages], stromal cells, and TCs). The scoring scale was (in %) 0, 1, 2, 3, 4, 5, 10, and further increments of 10 up to 100. Samples with LAG-3–positive IC percentage scores of ≥ 1% were reported as LAG-3–positive. Slides were randomized prior to evaluation. For each staining platform, all 80 slides were scored over 2 days (40 slides/day) followed by a washout period of 5 days. All 320 slides were scored by the same pathologist.

### Digital scoring

Images of all consecutive sections from the same tissue block evaluated using BOND-RX 1, ASL-48, VBU or BOND-RX 2 were analyzed in Visiopharm (Visiopharm, Hoersholm, Denmark). The analysis was performed using a single in-house scoring algorithm modified for use on the staining platforms utilized in this study, with a simple thresholding change and minor clean-up steps to eliminate some pigmentation. To ensure that a consistent region of interest (ROI [the tumor region containing the tumor and tumor-associated stroma]) was captured for serial sections, the ROI was annotated on 1 image and copied onto the other 3 images, with minor adjustments to account for interslide variability. Areas that were unevaluable on 1 image (eg, due to tissue detachment or excess melanin pigmentation) were not taken into consideration on the other images. If a section stained on 1 platform was completely unevaluable it was excluded, the sections stained on the other 3 platforms were analyzed, and a comment was made. If all sections from a block were unevaluable, then none were analyzed, and a comment was made. The digital scoring algorithm recorded LAG-3 expression as the sum of the positive 3,3’-diaminobenzidine tetrahydrochloride hydrate (DAB) signal as a fraction of the area of analysis.

### Statistical analysis

LAG-3 IHC scores were analyzed on the ratio-scale, and no additional normalization and/or standardization were applied. All analyses were performed using R software (version 4.0.5).

#### Analysis of LAG-3 using a numerical score

For the bar plot of LAG-3 score by case, cases were ordered from lowest to highest based on the values from the BOND-RX 2 samples. The prevalence as a function of cutoff was calculated by counting the percentage of cases exceeding (greater than or equal to) the given cutoff. For the scatter plot analyses of the manual and digital scoring data, the square-root transformation was applied to both X and Y axes for better visualization in the low-score range. For each scatter plot comparison, the strength of correlation was assessed using the coefficient of determination (*R*^*2*^) and Pearson correlation.

#### Analysis of LAG-3 positivity using a ≥ 1% cutoff

Agreement between platforms with a ≥ 1% cutoff to determine LAG-3 positivity was assessed using the pair-wise percentage agreement as well as Cohen’s Kappa per FDA guidelines and as previously described.^12, 13^ For this analysis, (i) in “A versus B percentage agreement”, A was used as the reference, (ii) the discordance metric was calculated as (100−overall percentage agreement)%, (iii) the 95% confidence interval (CI) for the percentage agreements were calculated using the Clopper–Pearson method,^14^ and (iv) the 95% CI for the Cohen’s Kappa was calculated using the variance estimates by Fleiss–Cohen–Everitt method.^15^ The Venn diagram was generated using the R package VennDiagram (version 1.6.0).

## Results

After adapting the BOND-RX staining procedure for use on ASL-48 and VBU, we examined the similarity of LAG-3 IHC staining patterns and intensities across the different staining platforms using consecutive sections cut from melanoma FFPE tissue blocks as shown in **Figure 1**. LAG-3 staining was assessed by visual inspection by a pathologist and revealed similar LAG-3 IHC staining patterns and intensities across the different staining platforms (**Figure 2**).

**Figure 2.**
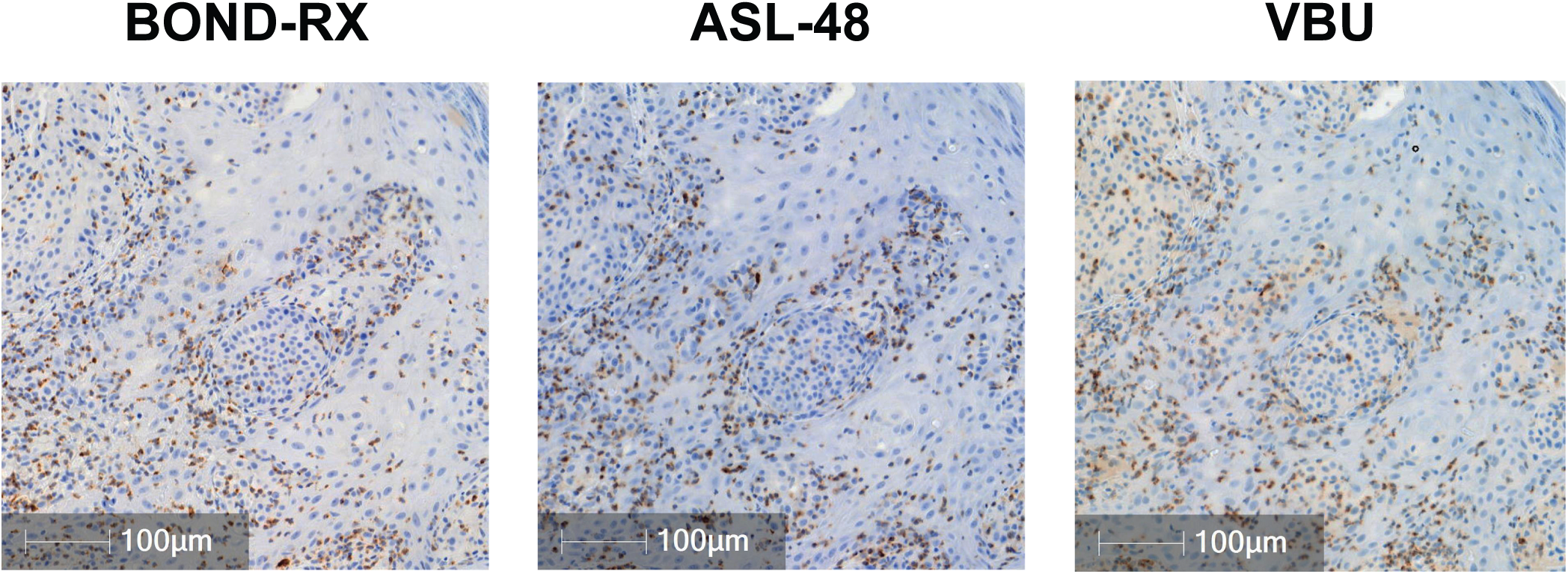
Visual comparison of LAG-3 staining in melanoma samples across 3 staining platforms. Representative images of LAG-3 IC IHC staining of consecutive sections cut from a commercially procured human FFPE melanoma tissue block using 3 IHC staining platforms: BOND-RX (left), ASL-48 (center), VBU (right). ASL-48, Agilent/Dako Autostainer Link 48; BOND-RX, Leica BOND-RX; FFPE, formalin-fixed paraffin-embedded; IC, immune cell; IHC, immunohistochemistry; LAG-3, lymphocyte- activation gene 3; VBU, VENTANA BenchMark ULTRA.

Next, we sought to compare pathologist scoring of LAG-3 expressed on ICs using a numerical score across the 3 staining platforms. Consecutive sections were cut from 80 melanoma FFPE tissue blocks and stained for LAG-3. Overall, for all cases, LAG-3 expression on ICs was comparable between consecutive sections cut from the same tissue block when stained using the different IHC platforms (**Figure 3A**). Similarly, consecutive sections cut from the same tissue block displayed comparable LAG-3 expression on ICs when both sections were stained using the BOND-RX IHC platform. The prevalence of LAG-3 expression on ICs at different cutoffs was similar across platforms and between runs on the same platform for the 2 BOND-RX runs (**Figure 3B**). For example, the percentage of samples determined as having LAG-3 IC expression ≥ 1% was 42% (34/80) for BOND-RX 1, 48% (38/80) for ASL-48, 56% (45/80) for VBU, and 51% (41/80) for BOND-RX 2. LAG-3 IC expression was strongly correlated between the 2 BOND-RX runs and between BOND-RX and both ASL-48 and VBU (**Figure 4A**). Notably, intraplatform concordance (BOND-RX 2 versus BOND-RX 1: Pearson correlation = 0.91 [*P* < 2.2e-16], slope = 1.2) was comparable to interplatform concordance (ASL-48 versus BOND-RX 1: Pearson correlation = 0.90 [*P* < 2.2e-16], slope = 1.2; VBU versus BOND-RX 2: Pearson correlation = 0.88 [*P* < 2.2e-16], slope = 0.9).

**Figure 3.**
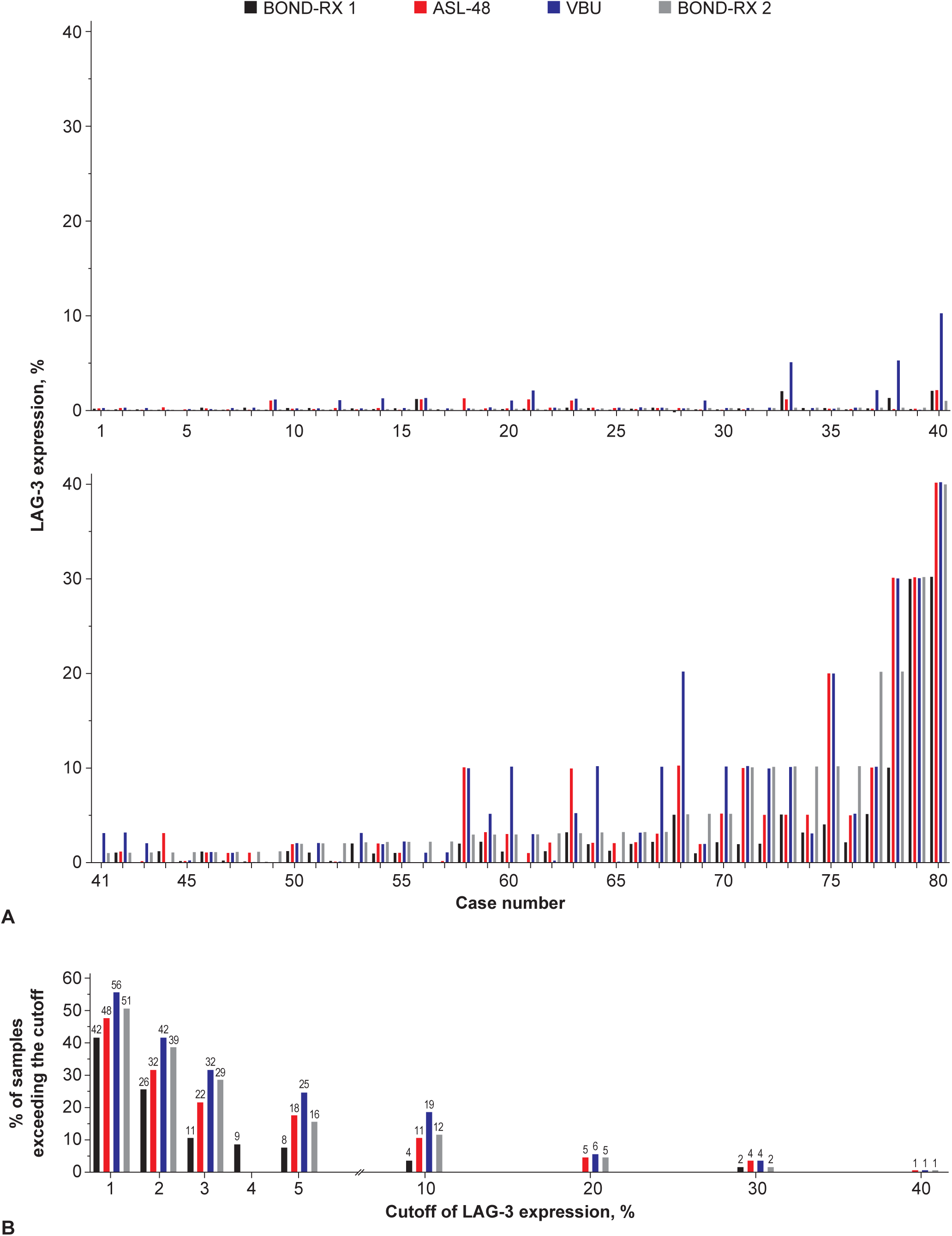
Comparison of LAG-3 IC IHC staining results across 3 staining platforms. **(A)** Bar charts show the IC expression of LAG-3 detected in consecutive sections cut from the same melanoma FFPE tissue block by the indicated staining platform. For the bar plot of LAG-3 score by case, cases were ordered from lowest to highest based on the values from the BOND-RX 2 samples. **(B)** Prevalence of LAG-3 IC expression by different cutoffs across different platforms. Bar charts show the percentage of samples exceeding (greater than or equal to) the indicated LAG-3 expression cutoffs. Two slides were run on the BOND-RX platform to assess intraplatform variability. ASL-48, Agilent/Dako Autostainer Link 48; BOND-RX 1, Leica BOND-RX [section 1]; BOND-RX 2, Leica BOND-RX [section 4]; FFPE, formalin-fixed paraffin-embedded; IC, immune cell; IHC, immunohistochemistry; LAG-3, lymphocyte-activation gene 3; VBU, VENTANA BenchMark ULTRA.

**Figure 4.**
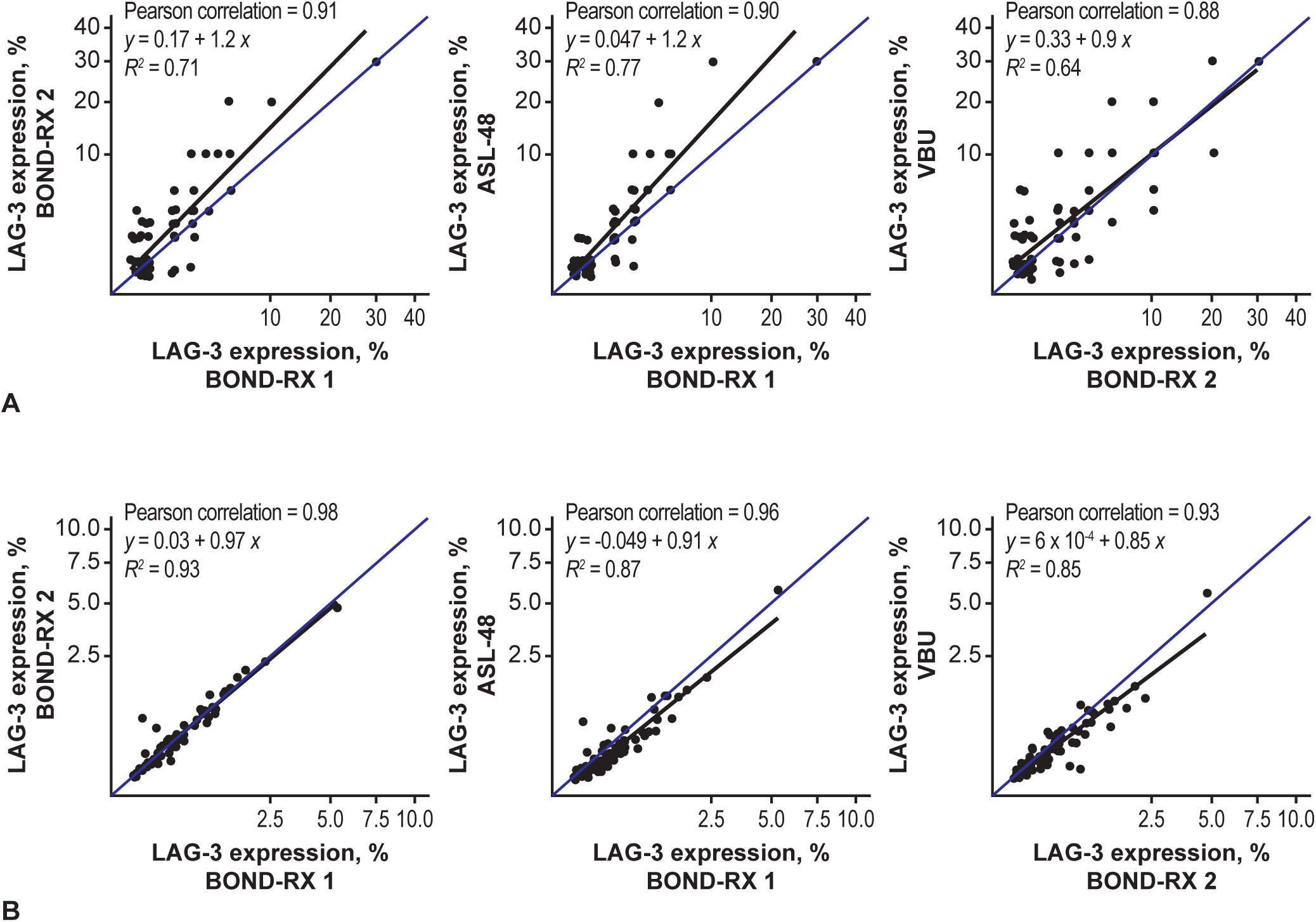
Scatter plots of platform comparison using pathologist scoring and a digital scoring algorithm. **(A)** Scatter plots of platform comparisons using pathologist scoring. LAG-3 expression was recorded using a numerical score defined as the percentage of LAG-3–positive ICs that morphologically resembled lymphocytes relative to all nucleated cells (ICs [lymphocytes and macrophages], stromal cells, and TCs). The blue line indicates the identity line, and the black line indicates the linear regression line. The coefficient of determination (*R*^2^) and Pearson correlation were calculated for each comparison and are displayed on the scatterplots. **(B)** Scatter plots of platform comparisons using the digital scoring algorithm. LAG-3 expression was recorded as the sum of the positive DAB signal as a fraction of the area of analysis. The blue line indicates the identity line, and the black line indicates the linear regression line. The coefficient of determination (*R*^2^) and Pearson correlation were calculated for each comparison and are displayed on the scatterplots. Two slides were run on the BOND- RX platform to assess intraplatform variability. ASL-48, Agilent/Dako Autostainer Link 48; BOND-RX 1, Leica BOND-RX [section 1]; BOND-RX 2, Leica BOND-RX [section 4]; DAB, 3,3’-diaminobenzidine tetrahydrochloride hydrate; IC, immune cell; LAG-3, lymphocyte-activation gene 3; TC, tumor cell; VBU, VENTANA BenchMark ULTRA.

Since observer variation is a documented limitation of quantitative IHC analysis by pathologists’ visual IC scoring, we also assessed LAG-3 expression by digital image analysis.^16-18^ This approach eliminates intraobserver variation and enables a more objective assessment of staining variability between platforms. Out of the 80 samples, LAG-3 expression was analyzed using the digital scoring algorithm in 77 samples on BOND-RX and ASL-48 and 75 samples on VBU. Although melanin was removed from all samples using the same method (**Table 1**), excess melanin pigmentation made 3 samples unevaluable by digital image analysis on all platforms, and a further 2 samples were unevaluable by digital image analysis on VBU. Notably, each of these samples had sufficient melanin removal for manual evaluation by a pathologist. Intraplatform and interplatform correlations increased when using the digital scoring algorithm compared with pathologist scoring (BOND-RX 2 versus BOND-RX 1: Pearson correlation = 0.98 [*P* < 2.2e-16], slope = 0.97; ASL-48 versus BOND-RX 1: Pearson correlation = 0.96 [*P* < 2.2e-16], slope = 0.91; VBU versus BOND-RX 2: Pearson correlation = 0.93 [*P* < 2.2e-16], slope = 0.85) (**Figure 4B**).

Finally, as clinical trials often focus on stratifying patients using a defined cutoff, we investigated interplatform and intraplatform agreement when LAG-3 IC positivity was determined from pathologist scores using a ≥ 1% cutoff. The level of agreement was high across all platforms, with point estimates for comparisons across all platforms > 75% for positive percentage agreement, negative percentage agreement, and overall percentage agreement, and a low level of discordance (< 25% across all platforms) (**Figure 5A**). Across all platforms, 27 out of 80 samples were determined as LAG-3– negative, and 26 out of 80 samples were determined as LAG-3–positive (**Figure 5B**).

**Figure 5.**
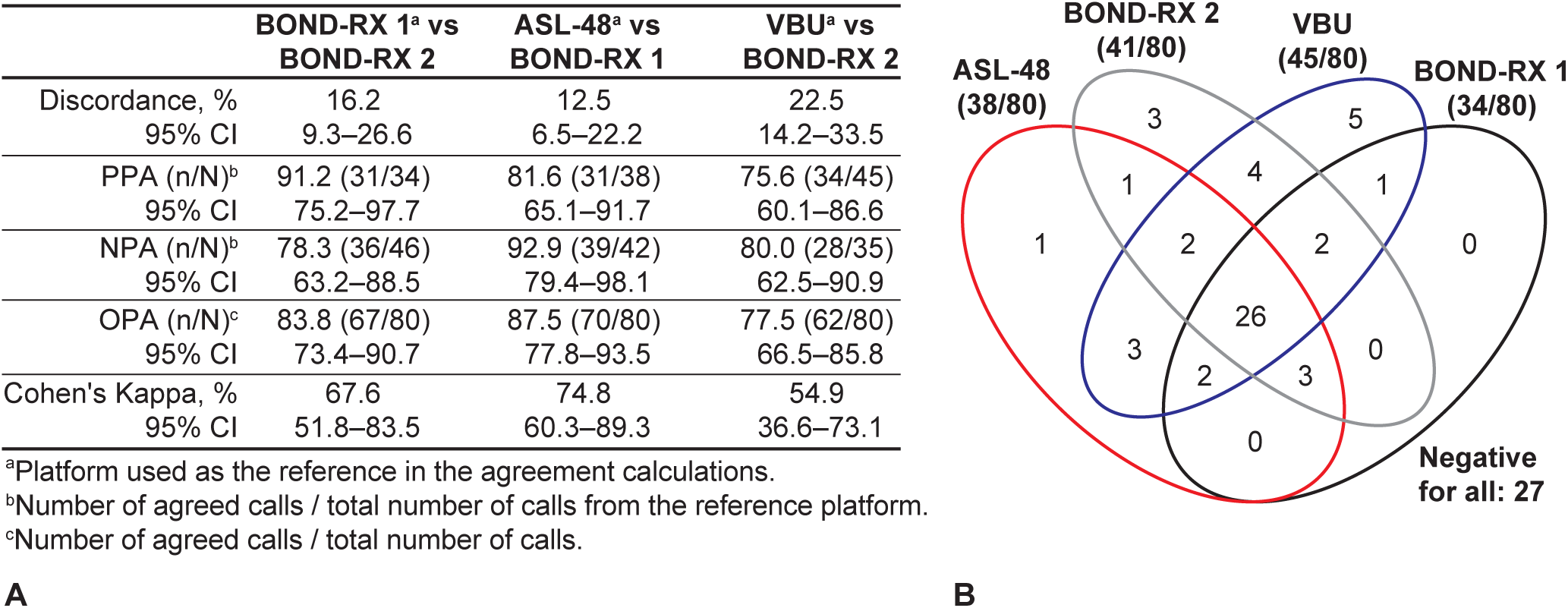
Agreement between 3 IHC staining platforms using a ≥ 1% cutoff to determine LAG-3 positivity. **(A)** Intraplatform and interplatform agreement between 3 IHC staining platforms. **(B)** A Venn diagram showing positive calls using a ≥ 1% cutoff to determine LAG-3 IC positivity across 3 IHC staining platforms. Two slides were run on the BOND-RX platform to assess intraplatform variability. ASL-48, Agilent/Dako Autostainer Link 48; BOND-RX 1, Leica BOND-RX [section 1]; BOND-RX 2, Leica BOND-RX [section 4]; CI, confidence interval; IC, immune cell; IHC, immunohistochemistry; LAG-3, lymphocyte-activation gene 3; NPA, negative percentage agreement; OPA, overall percentage agreement; PPA, positive percentage agreement; VBU, VENTANA BenchMark ULTRA.

## Discussion

This study investigated the comparability of IHC assays using the 17B4 antibody clone to assess LAG-3 expression across different staining platforms, which is an important consideration for the broad implementation of an IHC assay. Although the data from RELATIVITY-047 demonstrated that combined treatment with relatlimab and nivolumab prolongs PFS compared with nivolumab monotherapy regardless of LAG-3 status in patients with previously untreated metastatic or unresectable melanoma, there remains significant interest in assessing LAG-3 expression by IHC among the research community.^5^

Here, we show that LAG-3 IHC assays using the 17B4 antibody clone produce visually similar staining patterns and intensities across the BOND-RX, ASL-48, and VBU staining platforms. Comparable scores were observed across all 3 platforms when LAG-3 expression on ICs was measured by a pathologist using a numerical score. Pathologist agreement between platforms was similar when a ≥ 1% cutoff was used to determine LAG-3 positivity. Furthermore, pathologist agreement was consistent with previous publications and was in line with what would be expected in clinical settings.(Johnson et al, in preparation) As discussed in previous studies that reported comparable difficulties, the variance observed with pathologist scoring is likely due to issues associated with manually scoring IHC staining on ICs.^17, 18^ This task is more challenging than TC scoring due to IC size and variability. Importantly, intraplatform correlation and interplatform correlation were similar, highlighting the concordance of the LAG-3 IHC assays using the 17B4 antibody between staining platforms when performed manually by a pathologist.

In recent years, digital pathology techniques have been developed to address some of the inherent challenges within traditional manual pathology methods. In particular, digital pathology techniques are less time consuming and reduce interobserver variability compared with manual pathology methods.^16, 19^ In this study, the use of a digital scoring algorithm to analyze LAG-3 IC expression increased both intraplatform and interplatform correlation compared with pathologist scoring. This indicates that there is little true intraplatform and interplatform variance in LAG-3 IC staining. A challenge with digital scoring is that simple algorithms focused on color detection are susceptible to artifacts, due to a low signal-to-noise ratio caused by melanin presence.^16^ Improvements to digital scoring could be facilitated by further improvements in the removal of melanin, better color separation between the IHC chromogen and brown melanin pigment, or by developing artificial intelligence–based image analysis algorithms.

Previous IHC comparison studies have shown that specific antibodies can be used on alternative platforms in order to overcome barriers to implementation of IHC assays.^20-23^ For example, Hendry et al found that the 22C3 assay produced consistent programmed death ligand 1 (PD-L1) scoring on the ASL-48 platform and the VBU platform.^20^ Data presented here describe the development of LAG-3 IHC assays using the 17B4 antibody clone that produce comparable LAG-3 staining on the BOND-RX, VBU, and ASL-48 staining platforms.

An important limitation is that LAG-3 positivity determined using a ≥ 1% cutoff is not predictive of response to relatlimab, and therefore assessment of LAG-3 expression is for exploratory use only.^5^ Additionally, this study stained all samples in a single laboratory with one pathologist performing the scoring and did not assess interobserver variability within and across IHC staining platforms, or interantibody clone variability, as has been done in harmonization studies of PD-L1 IHC assays.^20, 22^ Although this ensured accurate assessment of interplatform reproducibility by preventing interlaboratory, interobserver, and interantibody clone bias, it does not reflect the variability that may be observed in real-world use of the assays. Notably, recently published data have demonstrated robust interobserver and interlaboratory reproducibility of the LAG-3 IHC assay using the 17B4 antibody clone on the Leica BOND-III staining platform.(Johnson et al., in preparation) Nevertheless, future studies examining interlaboratory, interobserver, and interantibody clone variability within and across additional platforms would provide further valuable evidence that LAG-3 IHC assays using the 17B4 antibody are reliable and suitable for use across a range of different staining platforms.

Here, we demonstrate that LAG-3 17B4-based IHC assays perform reproducibly across 3 different and widely used IHC staining platforms. Together with previously published data demonstrating the analytical precision and interlaboratory reproducibility of the LAG-3 17B4-based IHC assay on the Leica BOND-III (Johnson et al, in preparation), results presented here show that this assay is robust and could help overcome barriers to LAG-3 testing implementation in future studies.

## Authors’ Contributions

JW and JB designed the study. JW, KD, LD, and JB reviewed and interpreted the data. KA scored all study samples. AV developed the digital scoring algorithm and performed the digital image analysis. GG and CR developed and optimized the LAG-3 staining procedures on all IHC staining platforms. JB and CR oversaw the study.

## Acknowledgments

The authors acknowledge Iryna Shnitsar, PhD, of Bristol Myers Squibb for helpful discussions and contributions to the preparation of the manuscript. Medical writing and editorial support were provided by Peter Harrison, PhD, and Matthew Weddig of Spark Medica Inc, funded by Bristol Myers Squibb.

---

**Table 1.**
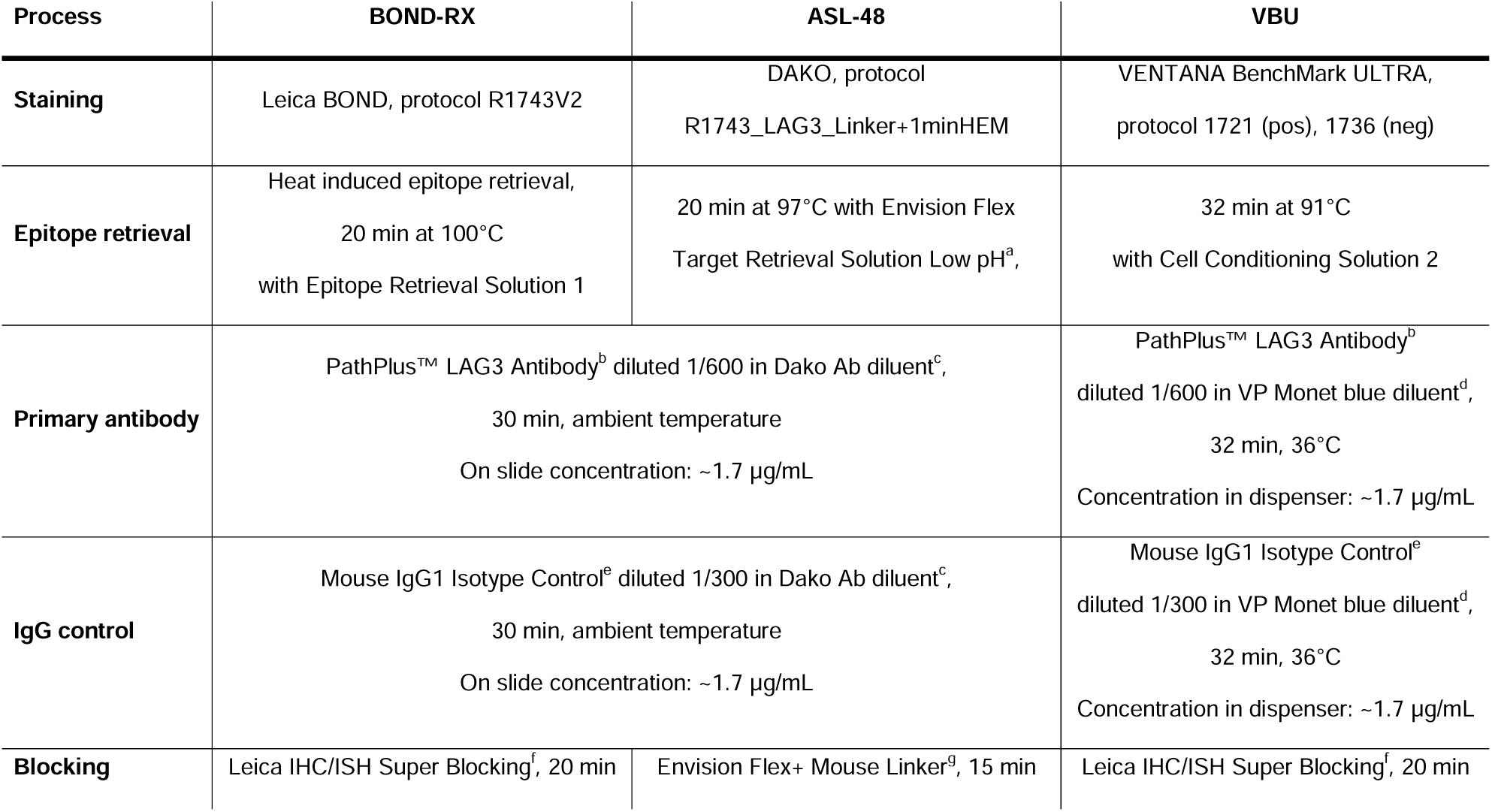
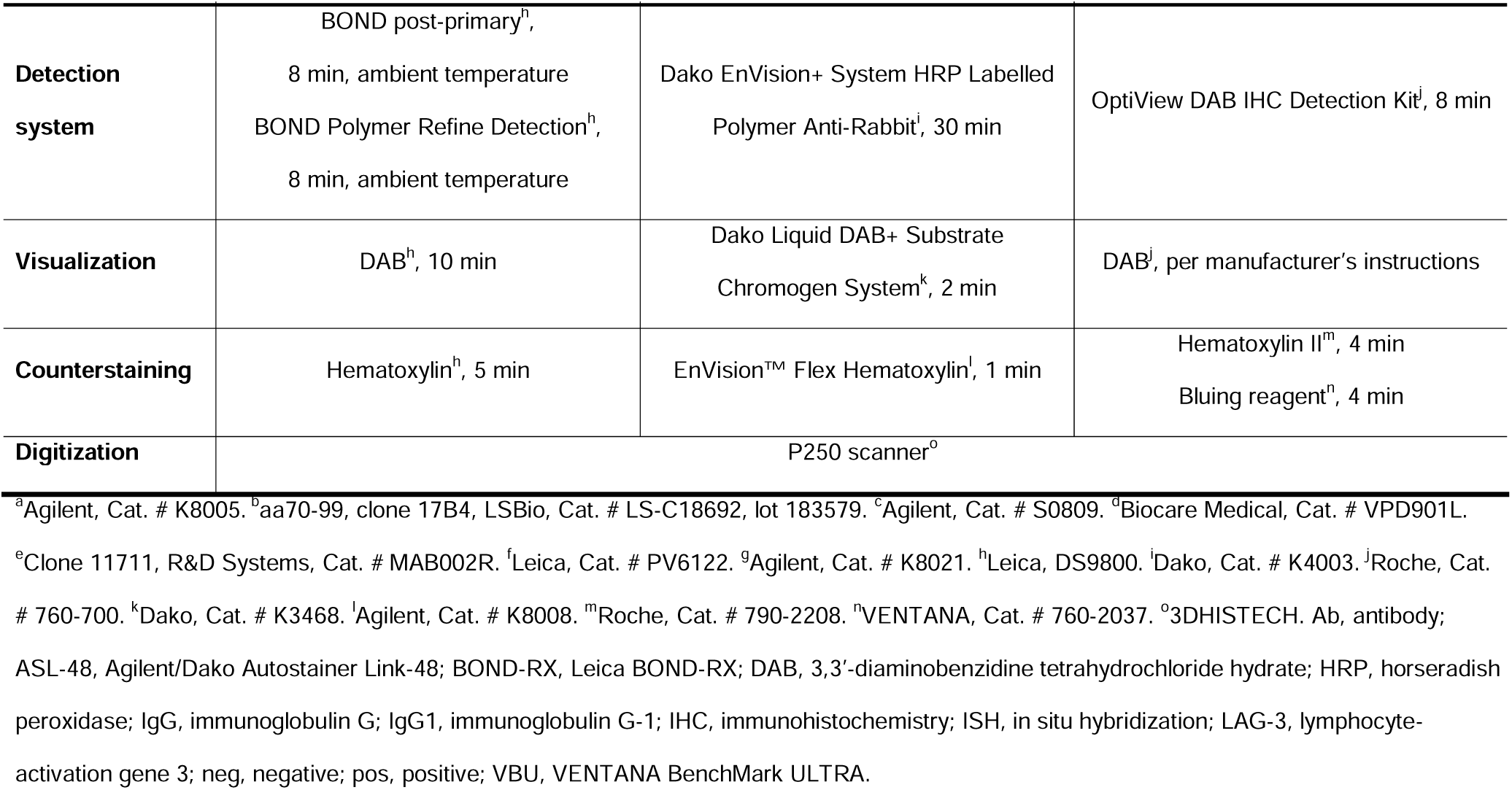
LAG-3 Staining Procedures on the Different IHC Staining Platforms Performed in a Single Laboratory.

